# Structural insights into Rad18 targeting by the SLF1 BRCT domains

**DOI:** 10.1101/2023.06.23.546354

**Authors:** Wei Huang, Fangjie Qiu, Lin Zheng, Meng Shi, Miaomiao Shen, Xiaolan Zhao, Song Xiang

**Author notes:** Correspondence for SX. These authors contributed equally to this work (Wei Huang, Fangjie Qiu).

## Abstract

Rad18 interacts with the SMC5/6 localization factor 1 (SLF1) to recruit the SMC5/6 complex to DNA damage sites for repair. The mechanism of the specific Rad18 recognition by SLF1 is unclear. Here, we present crystal structure of the tandem BRCT repeat (tBRCT) in SLF1 (SLF1^tBRCT^) bound with the interacting Rad18 peptide. Our structure and biochemical studies demonstrate that SLF1^tBRCT^ interacts with two phosphoserines and adjacent residues in Rad18 for high affinity and specificity Rad18 recognition. We found that SLF1^tBRCT^ utilizes mechanisms common among tBRCTs as well as unique ones for Rad18 binding, the latter include interactions with an α-helical structure in Rad18 that has not been observed in other tBRCT-bound ligand proteins. Our work provides structural insights into Rad18 targeting by SLF1 and expands the understanding of BRCT-mediated complex assembly.

## Introduction

The conserved Rad18 protein plays multiple roles in maintaining the genome stability. Its best-known function is to mono-ubiquitinate the proliferating cell nuclear antigen (PCNA), which signals for translesion DNA synthesis and enables PCNA poly-ubiquitination for additional DNA damage tolerance responses(1-4). Recently, Rad18 has been shown to interact with the structural maintenance of chromosome (SMC)5/6 localization factor 1 (SLF1)(5,6). The SMC5/6 complex has important functions in DNA repair(7,8). It was found that through interacting with SLF1, Rad18 recruits the SMC5/6 complex to DNA damage sites for repair(9,10). The importance of the Rad18-SLF1 pathway is highlighted by the fact that its deficiencies are associated with chromosomal instability and developmental defects(11).

Previous studies have shown that the tandem BRCA1 C-Terminal (BRCT) domain repeat (tBRCT) at the SLF1 N-terminus (SLF1^tBRCT^) plays a critical role in the Rad18-SLF1 association(5,6). BRCT domains were first identified in the tumor suppressor BRCA1 protein and subsequently found in more than twenty human proteins, most are involved in genome maintenance(12,13). In most cases, BRCT domains are assembled as tBRCT units, though units with single or more than two BRCT domains have also been found. Many tBRCTs specifically interact with phosphorylated ligand proteins, which often arise during DNA replication, DNA damage repair and checkpoint response. Such phosphorylation-dependent protein-protein interaction play key structural roles in the assembly of protein complexes in these processes as well as in the recruitment of proteins to specific genomic sites(14-17).

Structural studies revealed that the specificity of tBRCT-mediated ligand recognition is determined by both a phosphorylated serine or threonine residue and adjacent residues in the ligand proteins. The phosphorylation of Ser442 and Ser444 in Rad18 has been shown to be critical for the Rad18-SLF1 association(5). However, the adjacent Rad18 region shares no obvious sequence similarities with previously characterized tBRCT ligand proteins (Table S1). The molecular basis for the specific Rad18 targeting by SLF1 has been unclear.

Here, we demonstrate that SLF1^tBRCT^ interacts strongly and specifically with the Ser442/Ser444-containing Rad18 peptide upon phosphorylation of these serine residues. We determined the crystal structure of SLF1^tBRCT^ with bound Rad18 peptide. Structure-guided functional studies revealed that SLF1^tBRCT^ interacts with both phosphoserines as well as adjacent regions. Similar to other tBRCTs, two pockets in SLF1^tBRCT^ recognize phosphoserine 442 and a neighboring residue in Rad18. In addition, SLF1^tBRCT^ also forms three sets of interactions with Rad18 that diverge from interactions observed in other tBRCT-ligand complexes. Our study indicates that SLF1^tBRCT^ interacts with multiple residues in the Rad18 Ser442/Ser444-containing region to target Rad18 with high affinity and specificity.

## Results

### SLF1^tBRCT^ binds strongly and specifically to phosphoserine 442/444 containing Rad18 peptide

SLF1^tBRCT^ (residues 1-199) and the Rad18 Ser442/Ser444-containing region (residues 436-452) are conserved in mammals(5), birds and reptiles (Figs 1A and S1A-B). To probe the role of these regions in the Rad18-SLF1 association, we synthesized peptides corresponding to the Rad18 Ser442/Ser444-containing region and probed their interaction with SLF1^tBRCT^. Isothermal titration calorimetry (ITC) experiments indicated that SLF1^tBRCT^ interacts strongly with a Rad18 peptide containing phosphorylated Ser442 and Ser444 (Rad18-2P), with a *K*_D_ of 12 nM (Fig. 1B and Table 1). In contrast, its binding to an unphosphorylated Rad18 peptide (Rad18-NP) was undetectable by ITC.

**Figure 1.**
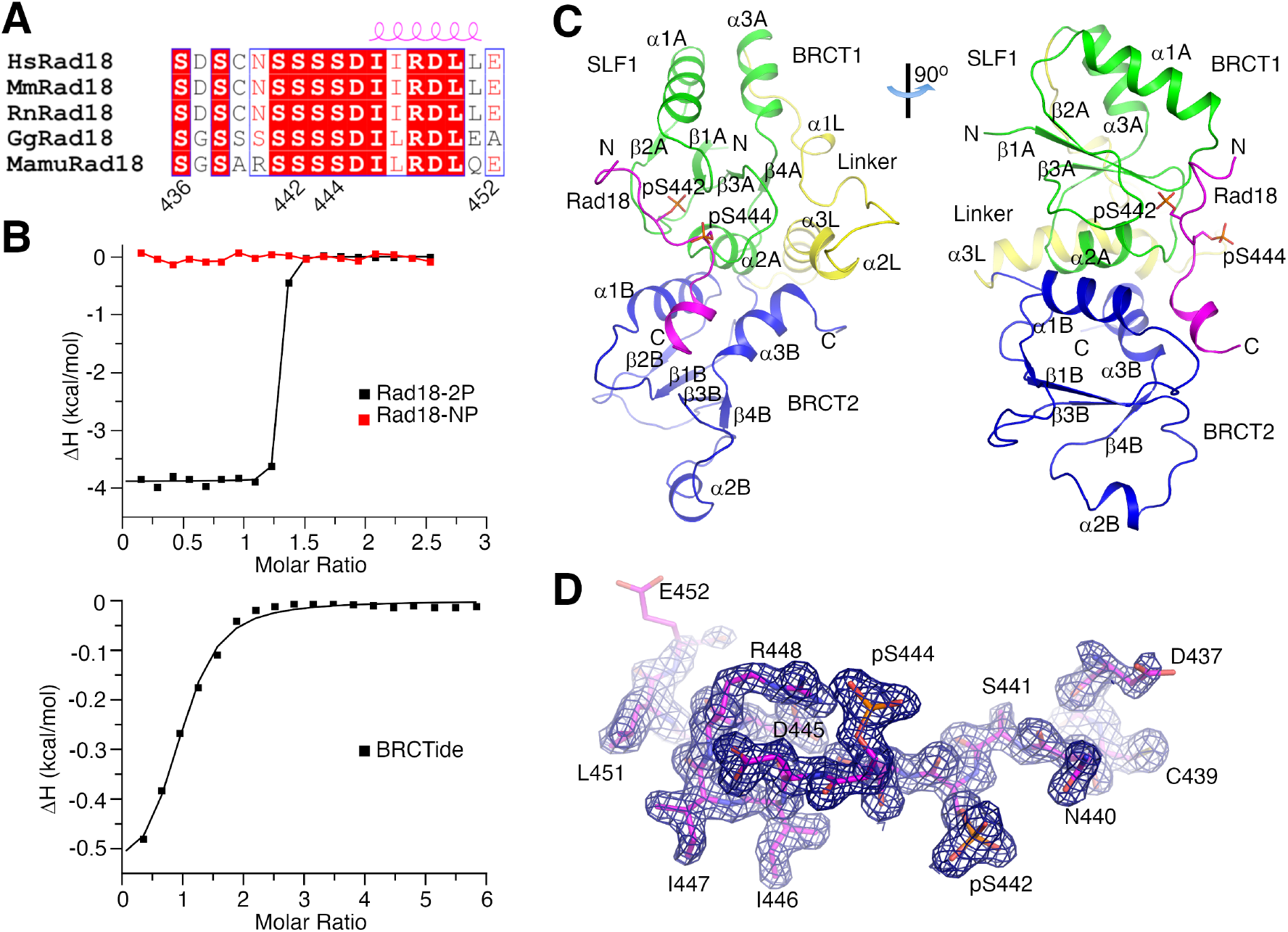
Structure of the SLF1^tBRCT^-Rad18-2P complex. (A) Sequence alignment of Rad18. Residue numbers and secondary structure elements for the human Rad18 are indicated. Hs, *Homo sapiens* (human); Mm, *Mus musculus* (mouse); Rn, *Rattus norvegicus* (rat); Gg, *Gallus gallus* (chicken); Mamu, *Mauremys mutica* (turtle). (B) ITC experiments probing the interaction between SLF1^tBRCT^ and Rad18 peptides or BRCTide. (C) Crystal structure of the SLF1^tBRCT^-Rad18-2P complex. SLF1 BRCT domains 1, 2 and the linker are colored in green, blue and yellow, respectively. Rad18 is colored in magenta. This color scheme is used throughout the manuscript unless otherwise indicated. Secondary structure elements and the N-and C-terminus of SLF1^tBRCT^ and the Rad18 peptide are indicated. (D) Electron density for the Rad18-2P peptide. The density for the peptide bound to SLF1^tBRCT^ molecule 1 in the crystal contoured at 1σ is presented. Structure figures are prepared with pymol (www.pymol.org).

**Table 1.**
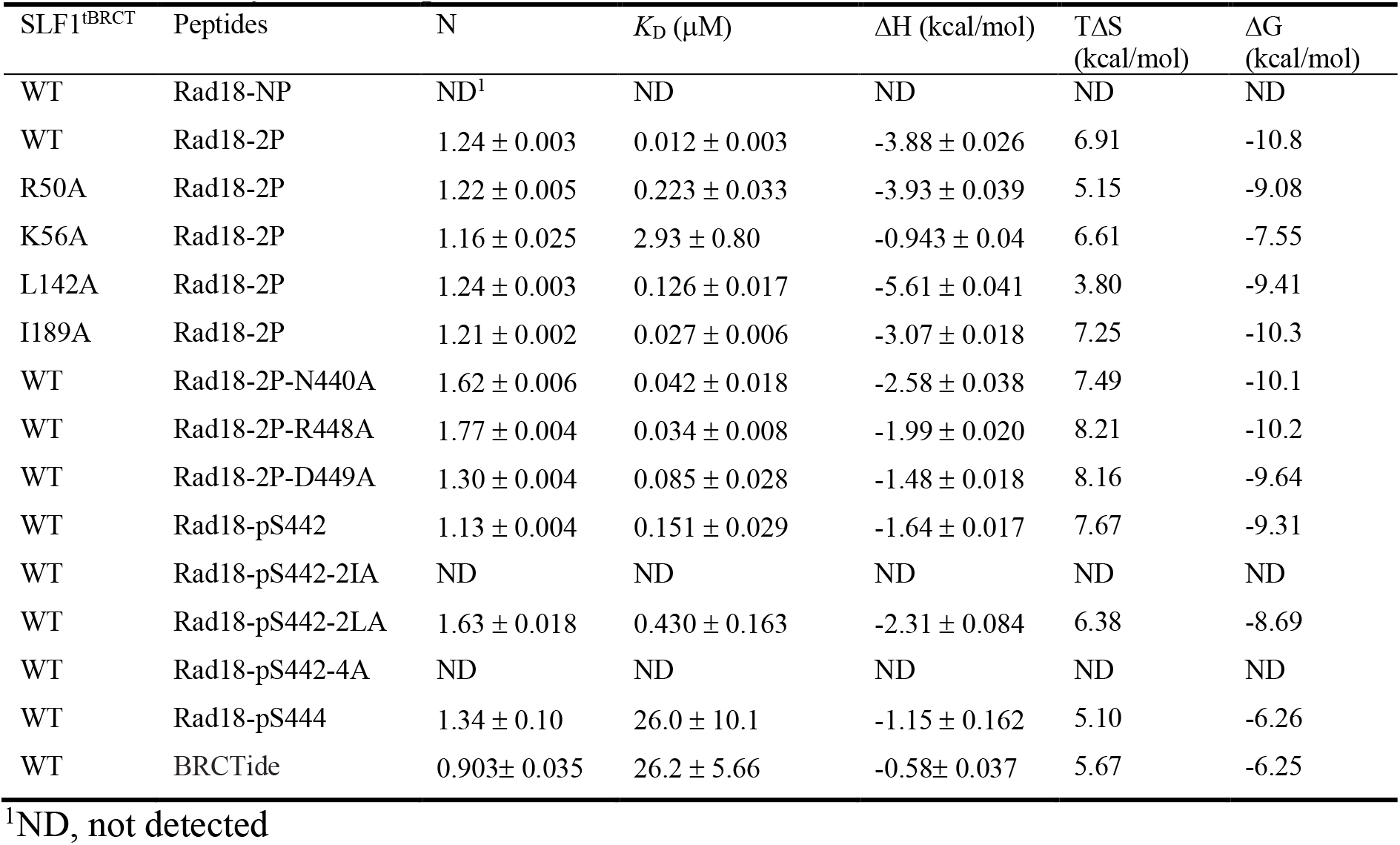
Summary of ITC experiments

| SLF1<sup>tBRCT</sup> | Peptides | N | $K_D$ (μM) | $\Delta H$ (kcal/mol) | $T\Delta S$ (kcal/mol) | $\Delta G$ (kcal/mol) |
| --- | --- | --- | --- | --- | --- | --- |
| WT | Rad18-NP | ND<sup>1</sup> | ND | ND | ND | ND |
| WT | Rad18-2P | $1.24 \pm 0.003$ | $0.012 \pm 0.003$ | $-3.88 \pm 0.026$ | 6.91 | -10.8 |
| R50A | Rad18-2P | $1.22 \pm 0.005$ | $0.223 \pm 0.033$ | $-3.93 \pm 0.039$ | 5.15 | -9.08 |
| K56A | Rad18-2P | $1.16 \pm 0.025$ | $2.93 \pm 0.80$ | $-0.943 \pm 0.04$ | 6.61 | -7.55 |
| L142A | Rad18-2P | $1.24 \pm 0.003$ | $0.126 \pm 0.017$ | $-5.61 \pm 0.041$ | 3.80 | -9.41 |
| I189A | Rad18-2P | $1.21 \pm 0.002$ | $0.027 \pm 0.006$ | $-3.07 \pm 0.018$ | 7.25 | -10.3 |
| WT | Rad18-2P-N440A | $1.62 \pm 0.006$ | $0.042 \pm 0.018$ | $-2.58 \pm 0.038$ | 7.49 | -10.1 |
| WT | Rad18-2P-R448A | $1.77 \pm 0.004$ | $0.034 \pm 0.008$ | $-1.99 \pm 0.020$ | 8.21 | -10.2 |
| WT | Rad18-2P-D449A | $1.30 \pm 0.004$ | $0.085 \pm 0.028$ | $-1.48 \pm 0.018$ | 8.16 | -9.64 |
| WT | Rad18-pS442 | $1.13 \pm 0.004$ | $0.151 \pm 0.029$ | $-1.64 \pm 0.017$ | 7.67 | -9.31 |
| WT | Rad18-pS442-2IA | ND | ND | ND | ND | ND |
| WT | Rad18-pS442-2LA | $1.63 \pm 0.018$ | $0.430 \pm 0.163$ | $-2.31 \pm 0.084$ | 6.38 | -8.69 |
| WT | Rad18-pS442-4A | ND | ND | ND | ND | ND |
| WT | Rad18-pS444 | $1.34 \pm 0.10$ | $26.0 \pm 10.1$ | $-1.15 \pm 0.162$ | 5.10 | -6.26 |
| WT | BRCTide | $0.903 \pm 0.035$ | $26.2 \pm 5.66$ | $-0.58 \pm 0.037$ | 5.67 | -6.25 |

<sup>1</sup>ND, not detected

To assess the specificity of the SLF1^tBRCT^-Rad18 interaction, we probed the interaction of SLF1^tBRCT^ with a phosphopeptide optimized for binding to the BRCA1 tBRCT (BRCTide, GAAYDI(pS)QVFPFAKKK)(18). ITC experiments indicated that SLF1^tBRCT^ interacted weakly with BRCTide, with a *K*_D_ of 26.2 μM that is ∼2000-fold higher than the *K*_D_ for the Rad18-2P peptide (Fig. 1B and Table 1). Collectively, our data suggest that Ser442 and Ser444 phosphorylation enables strong and specific interaction between the Rad18 Ser442/Ser444-containing region and SLF1^tBRCT^.

### Structure of SLF1^tBRCT^ in complex with a dually phosphorylated Rad18 peptide

The strong and specific Rad18-2P-SLF1^tBRCT^ interaction enabled us to crystalize their complex and determined the structure to a resolution of 1.75 Å (Table 2). SLF1 residues 5-199 are resolved in the electron density map, which form a compact and elongated structure with a dimension of 35×35×65 Å (Fig. 1C). The two SLF1 BRCT domains (BRCT 1 and 2) adopt typical BRCT fold, consisting of a central 4-strand parallel β-sheet flanked by two α-helices on one side and one on the other. They are arranged in parallel orientation and form primarily hydrophobic interactions with each other. The SLF1^tBRCT^ structure is further stabilized by the three-helix linker that interacts with both BRCT domains. Two SLF1^tBRCT^ molecules were found in the asymmetric unit. Most regions in them adopt very similar structures and can be aligned with a root mean square deviation (RMSD) of 0.69 Å for the Cα atoms (Fig. S2A). Structural differences were observed for part of the β1B-α1B loop (letters A and B indicate the first and second BRCT domains, respectively) and the α2B-spanning region, suggesting structural flexibility of these regions. Both regions participate in crystal packing interactions, which may stabilize the observed structure.

**Table 2.**
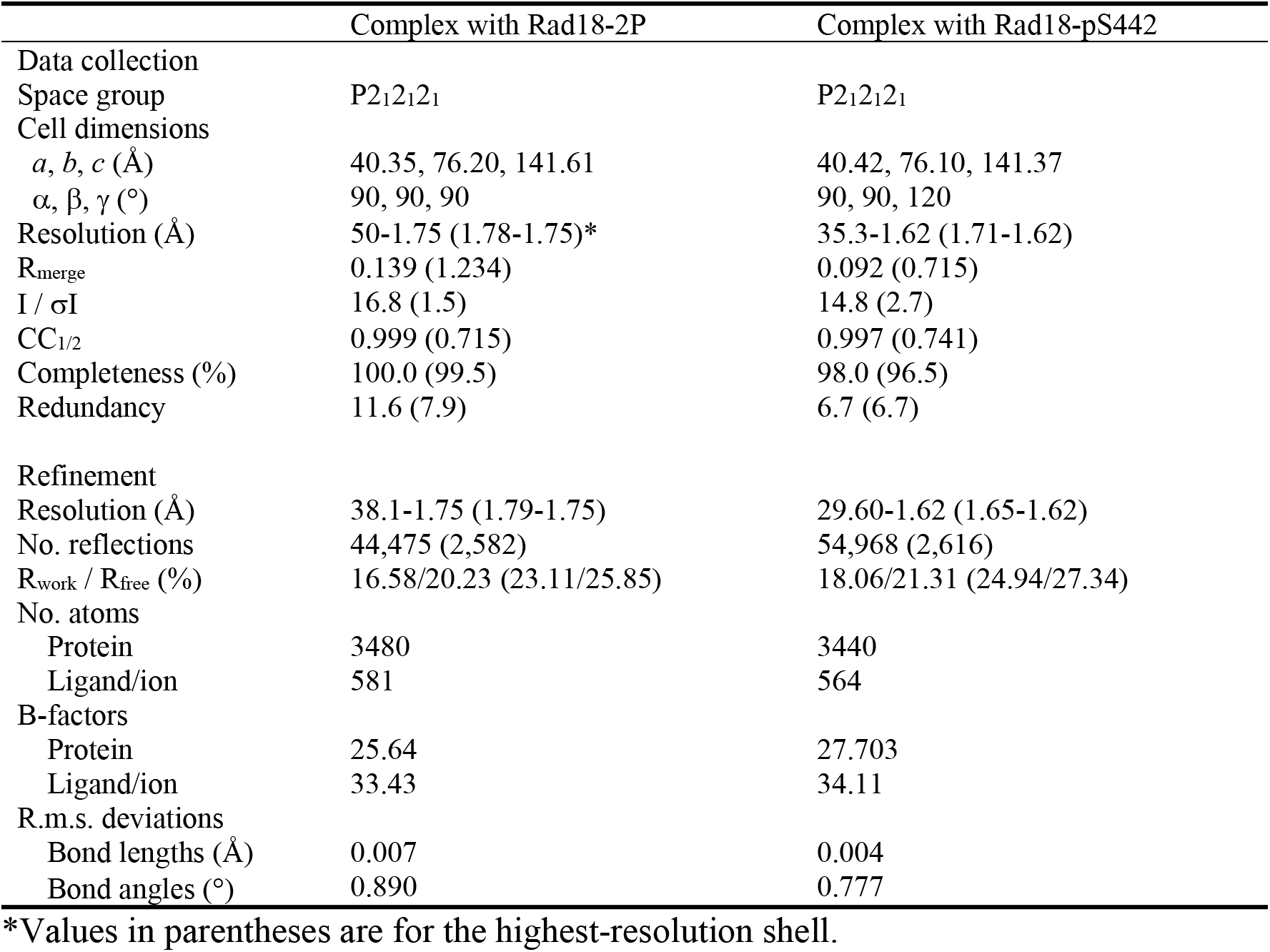
Data collection and structure refinement statistics

|  | Complex with Rad18-2P | Complex with Rad18-pS442 |
| --- | --- | --- |
| Data collection |  |  |
| Space group | P2<sub>1</sub>2<sub>1</sub>2<sub>1</sub> | P2<sub>1</sub>2<sub>1</sub>2<sub>1</sub> |
| Cell dimensions |  |  |
| *a*, *b*, *c* (Å) | 40.35, 76.20, 141.61 | 40.42, 76.10, 141.37 |
| $\alpha$, $\beta$, $\gamma$ (°) | 90, 90, 90 | 90, 90, 120 |
| Resolution (Å) | 50-1.75 (1.78-1.75)* | 35.3-1.62 (1.71-1.62) |
| R<sub>merge</sub> | 0.139 (1.234) | 0.092 (0.715) |
| I / σI | 16.8 (1.5) | 14.8 (2.7) |
| CC<sub>1/2</sub> | 0.999 (0.715) | 0.997 (0.741) |
| Completeness (%) | 100.0 (99.5) | 98.0 (96.5) |
| Redundancy | 11.6 (7.9) | 6.7 (6.7) |
| Refinement |  |  |
| Resolution (Å) | 38.1-1.75 (1.79-1.75) | 29.60-1.62 (1.65-1.62) |
| No. reflections | 44,475 (2,582) | 54,968 (2,616) |
| R<sub>work</sub> / R<sub>free</sub> (%) | 16.58/20.23 (23.11/25.85) | 18.06/21.31 (24.94/27.34) |
| No. atoms |  |  |
| Protein | 3480 | 3440 |
| Ligand/ion | 581 | 564 |
| B-factors |  |  |
| Protein | 25.64 | 27.703 |
| Ligand/ion | 33.43 | 34.11 |
| R.m.s. deviations |  |  |
| Bond lengths (Å) | 0.007 | 0.004 |
| Bond angles (°) | 0.890 | 0.777 |

\*Values in parentheses are for the highest-resolution shell.

Strong densities for the Rad18-2P peptide were found on both SLF1^tBRCT^ molecules in the crystal (Figs 1D and S2B). Rad18 residues 437-452 and 436-451 were fitted into these densities on SLF1^tBRCT^ molecules 1 and 2, respectively, revealing mostly identical structures for the two Rad18-2P peptides (Fig. S2A). The Rad18 binding site in SLF1^tBRCT^ is composed by both BRCT domains and predominantly positively charged (Fig. S2C). The Rad18 peptide N-terminal half including the phosphoserines adopt an extended conformation and interacts with BRCT 1, its C-terminal half forms an α-helix and interacts primarily with BRCT 2 (Fig. 1C). The Rad18-2P-SLF1^tBRCT^ interface is the largest among tBRCT-ligand peptide structures reported to date, burying 1600 Å^2^ of surface area.

### Phosphoserine 442 and 444 in Rad18 make distinct contributions to SLF1 binding

Our structure revealed that phosphoserines 442 and 444 (pS442 and pS444) in Rad18 form distinct interactions with SLF1^tBRCT^ (Fig. 2A). The pS442 side chain is completely buried at the Rad18-SLF1 interface. Its phosphate group forms polar interactions with the SLF1 Thr13 and Lys56 side chains and the Gly14 main chain carbonyl. In contrast, the pS444 side chain is mostly exposed to solvent. Its phosphate group forms polar interactions with the SLF1 Arg50 side chain in complex 2 in the crystal (Fig. S2D). It also interacts with the Arg448 side chain in Rad18 (Figs 2A and S2D).

**Figure 2.**
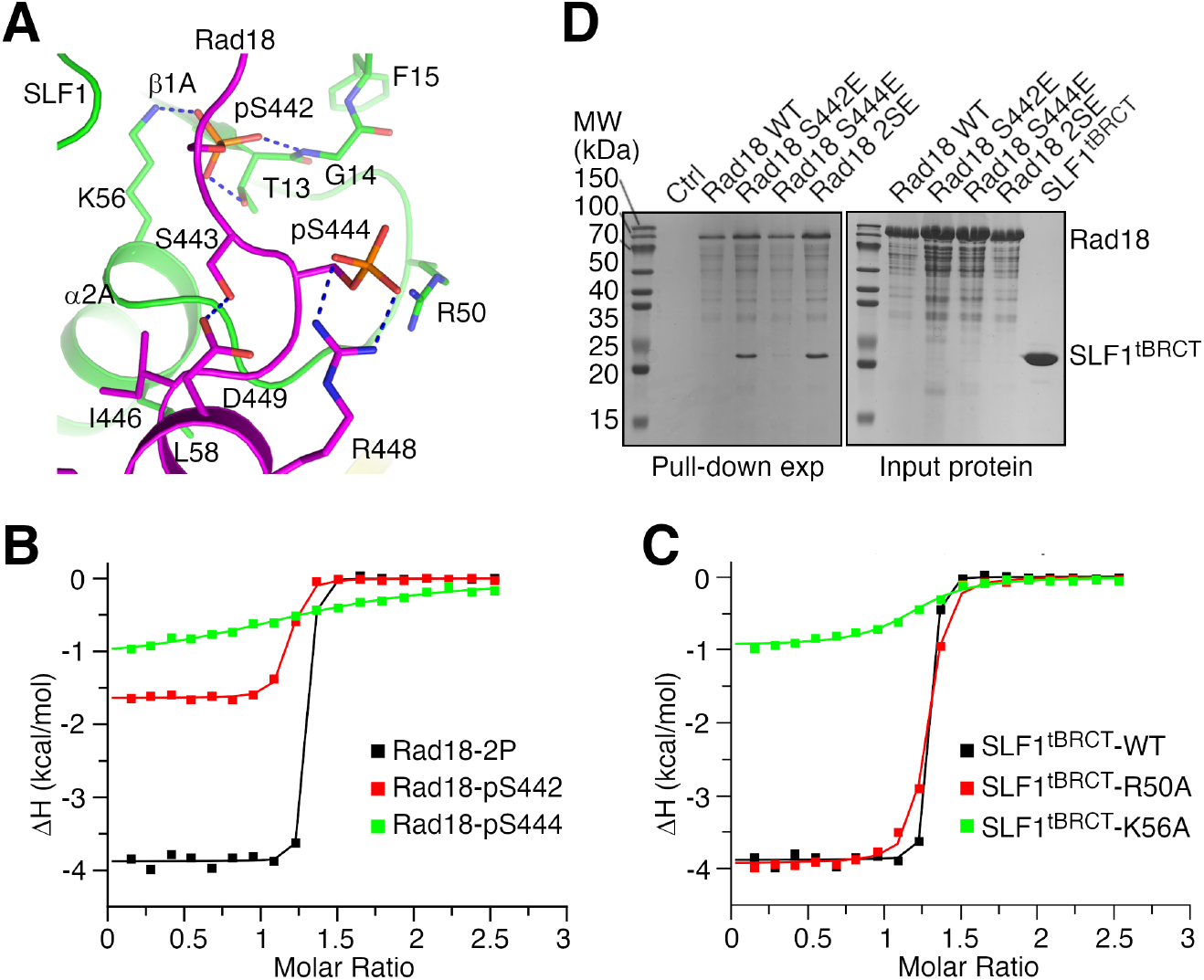
Phosphoserine 442 and 444 in Rad18 make distinct contributions to its interaction with SLF1^tBRCT^. (A) Interactions mediated by pS442, pS444 and surrounding regions in Rad18. Structure of complex 1 in the crystal is presented. Important residue side chains are highlighted. Dashed lines indicate hydrogen bonds or polar interactions. (B) ITC experiments probing the interaction between SLF1^tBRCT^ and Rad18 peptides. (C) ITC experiments probing the interaction between the Rad18-2P peptide and substituted SLF1^tBRCT^. Data for the wild type SLF1^tBRCT^ is included for comparison. (D) Rad18-SLF1^tBRCT^ pull-down experiments. Sodium dodecyl sulfate (SDS) polyacrylamide gel electrophoresis (PAGE) analysis of SLF1^tBRCT^ co-precipitated with the wild type or substituted Rad18 is presented. In lane “Ctrl”, SDS PAGE analysis of the experiment without Rad18 is presented.

To probe the function of pS442 and pS444 in the Rad18-SLF1^tBRCT^ association, we evaluated the effects of removing one of the phosphate groups or substituting SLF1 residues they interact with. ITC experiments revealed that removing the pS442 phosphate group or alanine substitution of its interacting residue Lys56 in SLF1^tBRCT^ strongly inhibited the Rad18-2P-SLF1^tBRCT^ interaction, resulting in 2100- and 240-fold increases in *K*_D_, respectively (Figs 2B-C and Table 1). In contrast, moderate effects were observed for removing the pS444 phosphate group or alanine substitution of its interacting residue Arg50. These modifications caused 13- and 19-fold increases in *K*_D_, respectively (Figs 2B-C and Table 1). These ITC data suggest that Ser442 phosphorylation is critical for the Rad18-SLF1^tBRCT^ association, whereas Ser444 phosphorylation plays a less important role and enhances the binding affinity.

To validate the ITC data, we carried out *in vitro* pull-down experiments with purified full-length Rad18 and SLF1^tBRCT^. Serine to glutamate substitutions were introduced into Rad18 to mimic the Serine 442 and/or 444 phosphorylation. The pull-down experiments indicated that SLF1^tBRCT^ strongly co-precipitated with the S442E- or S442E/S444E (2SE)-substituted Rad18, but not the wild type or the S444E-substituted Rad18 (Fig. 2D). These pull-down data are in line with the notion that Ser442 and Ser444 phosphorylation make different contributions to the Rad18-SLF1^tBRCT^ association. However, these data do not exclude a role of pS444 in promoting Rad18-SLF1^tBRCT^ association, as the serine to glutamate substitution does not fully replicate physiological functions of serine phosphorylation.

### Structure of SLF1^tBRCT^ in complex with a Rad18-pS442 peptide

Our binding experiments suggest that Ser442 phosphorylation is critical and sufficient for strong Rad18-SLF1^tBRCT^ association. To further clarify the roles of Ser442 and 444 phosphorylation in the Rad18-SLF1 interaction, we co-crystallized SLF1^tBRCT^ with the singularly phosphorylated Rad18-pS442 peptide and determined its structure to a resolution of 1.62 Å (Table 2). Two SLF1^tBRCT^ molecules were found in the asymmetric unit and electron densities for the Rad18-pSer442 peptide were found on both of them (Figs 3A and S3A). The structure and crystal packing of the SLF1^tBRCT^-Rad18-pS442 complex are very similar to the SLF1^tBRCT^-Rad18-2P complex (Fig. 3B). Compared to the SLF1^tBRCT^-Rad18-2P complex, little structural differences were observed even for the SLF1 Arg50 and Rad18 Arg448 side chains, which interact with the pS444 phosphate group in the Rad18-2P peptide (Figs 3C and S3B). The structure is consistent with the critical role of Ser442 phosphorylation in the Rad18-SLF1 association.

**Figure 3.**
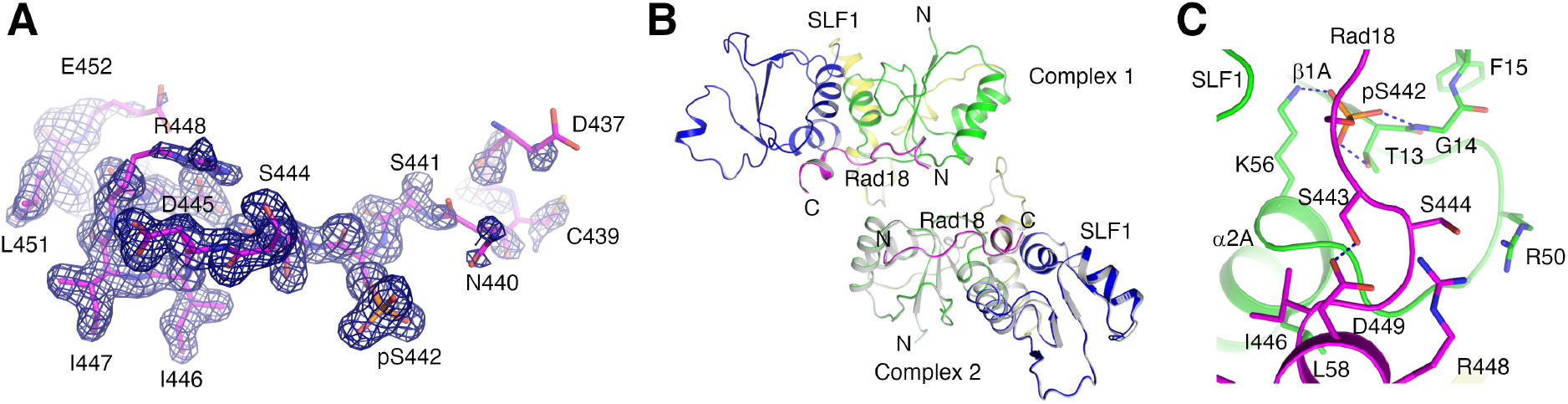
Structure of the SLF1^tBRCT^-Rad18-pS442 complex. (A) Electron density for the Rad18-pS442 peptide. The density for the Rad18 peptide bound to SLF1^tBRCT^ molecule 1 in the crystal contoured at 1σ is presented. (B) Comparison of the asymmetric unit structures. Superimposition of asymmetric unit structures of the SLF1^tBRCT^-Rad18-pS442 complex and the SLF1^tBRCT^-Rad18-2P complex (gray) is shown. The RMSD for the equivalent Cα atoms is 0.26 Å. (C) Interactions mediated by pS442 and surrounding regions in the Rad18-pS442 peptide. Structure of complex 1 in the crystal is presented.

The electron densities for the Rad18-pS442 peptide are weaker compared to electron densities for the Rad18-2P peptide, especially at the N-terminal region where Ser442 and Ser444 reside (Figs 3A and S3A). Such difference in density is in line with the reduced affinity of the Rad18-pS442 peptide towards SLF1^tBRCT^ and supports the function of Ser444 phosphorylation in enhancing the Rad18-SLF1^tBRCT^ binding. Electron densities for the Arg448 side chain in the Rad18-pS442 peptide is also weaker, consistent with the lack of Arg448-pS444 interaction in this peptide.

### Rad18 regions adjacent to Ser442 and Ser444 are critical for interaction with SLF1^tBRCT^

Our structures revealed that Rad18 residues adjacent to Ser442 and Ser444 form two additional interfaces with SLF1^tBRCT^. The Rad18-2P peptide N-terminal region forms several hydrogen bonds with SLF1^tBRCT^. Its Cys439 main chain carboxyl hydrogen bonds with the SLF1 Lys36 side chain, Asn440 side chain hydrogen bonds with the SLF1 Lys20 side chain and the Phe15 main chain amide (Figs 4A and S4A). Its C terminal half adopts an α-helical structure and forms an extensive interface with SLF1^tBRCT^, presenting residues Ile446, Ile447, Leu450 and Leu451 for hydrophobic interactions with SLF1 residues Leu58, Val133, Ser141, Leu142, Val145, Ile189, Leu192 and Gly193 (Figs 4B and S4B). The α-helix is stabilized by the pS444-Arg448 interaction and an Ser443-Asp449 hydrogen bond. The same regions in the Rad18-pS442 peptide form virtually identical interactions with SLF1^tBRCT^, although the pS444-Arg448 interaction is absent in this peptide.

**Figure 4.**
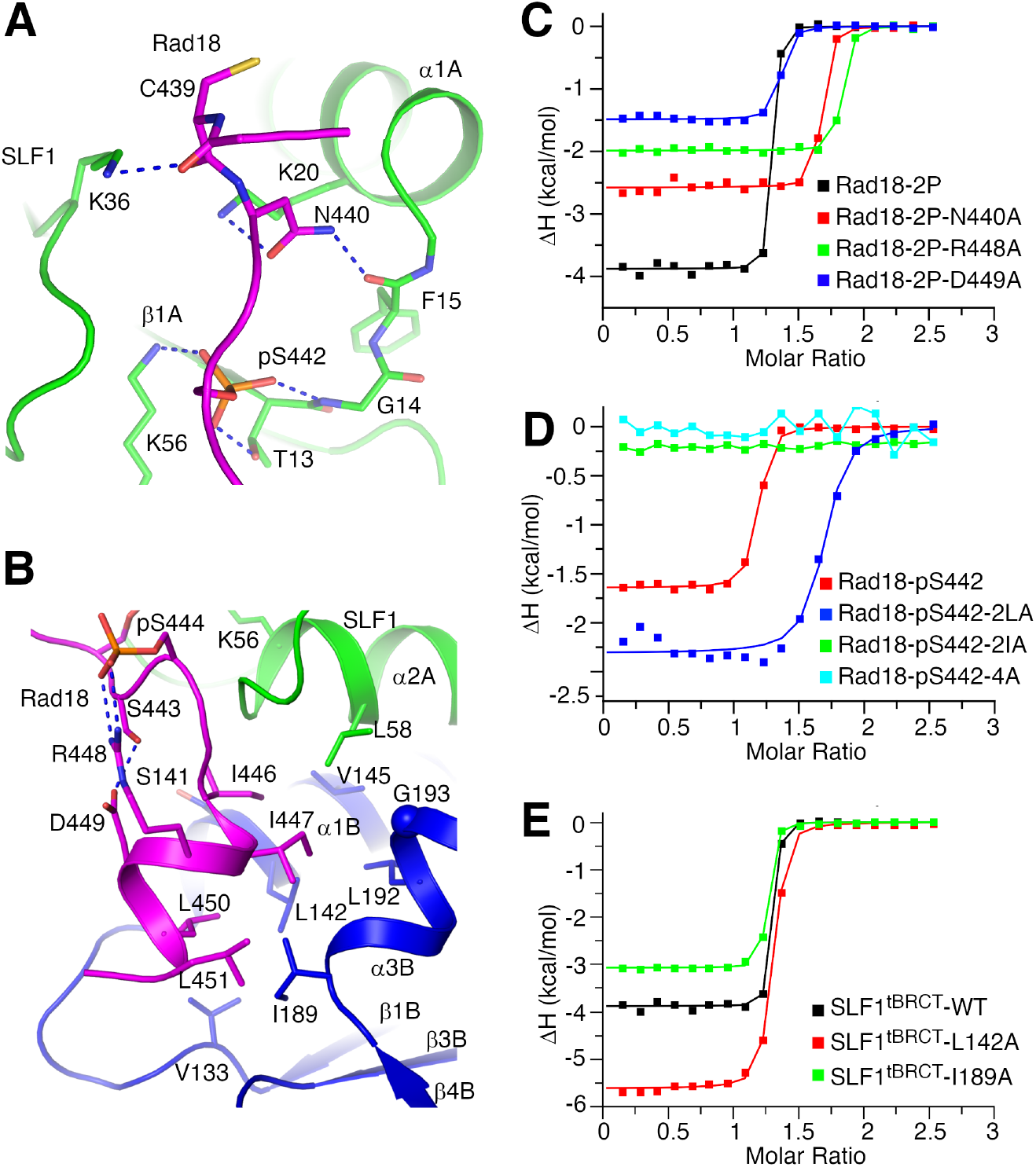
Rad18 regions adjacent to Ser442 and Ser444 make important contributions to its interaction with SLF1^tBRCT^. (A) and (B) interactions mediated by the N-(A) and C-terminal (B) regions of the Rad18-2P peptide. Structure of complex 1 in the crystal is presented. (C)-(D) ITC experiments probing the interaction between SLF1^tBRCT^ and substituted Rad18-2P (C) or Rad18-pS442 (D) peptides. Data for the wild type Rad18 peptides are included for comparison. (E) ITC experiments probing the interaction between the Rad18-2P peptide and substituted SLF1^tBRCT^. Data for the wild type SLF1^tBRCT^ is included for comparison.

To probe the function of the above mentioned SLF1-interacting residues in Rad18, we evaluated the effect of their substitutions on the Rad18-SLF1^tBRCT^ interaction. ITC experiments revealed moderate inhibitions of interaction for the N440A substitution at the Rad18 peptide N-terminus and the L450A/L451A (2LA) substitution at the C-terminal half of the Rad18 α-helix. These substitutions caused 3.5- and 2.8-fold increases in *K*_D_, respectively (Figs 4C-D and Table 1). In contrast, strong inhibitions were observed for the I446A/I447A (2IA) substitution at the N-terminal half of the Rad18 α-helix and its combination with 2LA (4A). Both substitutions reduced the Rad18-SLF1^tBRCT^ interaction to undetectable levels (Fig. 4D). To further probe the function of the Rad18 α-helix in binding SLF1^tBRCT^, we introduced the R448A and D449A substitutions in Rad18 to disrupt interactions stabilizing the α-helix, and alanine substitutions of Leu142 and Ile189 in SLF1^tRBRCT^ that interact with it. ITC experiments revealed that these substitutions increased the *K*_D_ 2-to 10-fold (Figs 4C and E and Table 1), supporting an important role of the Rad18 α-helix in its interaction with SLF1^tBRCT^. Interestingly, the L142A substitution in SLF1^tBRCT^ caused a much stronger inhibition than the I189A substitution. Leu142 and Ile189 interact primarily with the N-and C-terminal halves of the Rad18 α-helix, respectively. Such data is in line with the notion that the N-terminal half of the Rad18 α-helix plays a more important role than the C-terminal half in interacting with SLF1^tBRCT^.

Together, our structure and the ITC data suggest that Rad18 regions adjacent to Ser442 and Ser444 also contribute to the Rad18-SLF1^tBRCT^ interaction, among which the α-helix in Rad18 plays a particularly important role.

### Structural comparison with other tBRCT-ligand complexes

Tandem BRCT repeats can adopt several different conformations(18-22). Despite limited sequence similarity (Fig. 5A), SLF1^tBRCT^ is structurally homologous to ten other tBRCTs, including tBRCTs in the human BRCA1 (Fig. S5)(18,19) and BARD1(23), the fission yeast Brc1 (BRCT domains 5/6)(24) and its homologs Rtt107 in budding yeast (BRCT domains 5/6)(25) and PTIP in human (BRCT domains 5/6), the fission yeast Crb2(26) and its homolog 53BP1 in human(27), and three other human proteins, namely TopBP1 (BRCT domains 7/8)(28), MDC1(29,30), and MCPH1(31-33). Except for the BARD1 tBRCT, structures of these tBRCTs bound with phosphorylated ligand peptides have been reported, allowing comparative structural analysis to further understand the Rad18-SLF1^tBRCT^ interaction.

**Figure 5.**
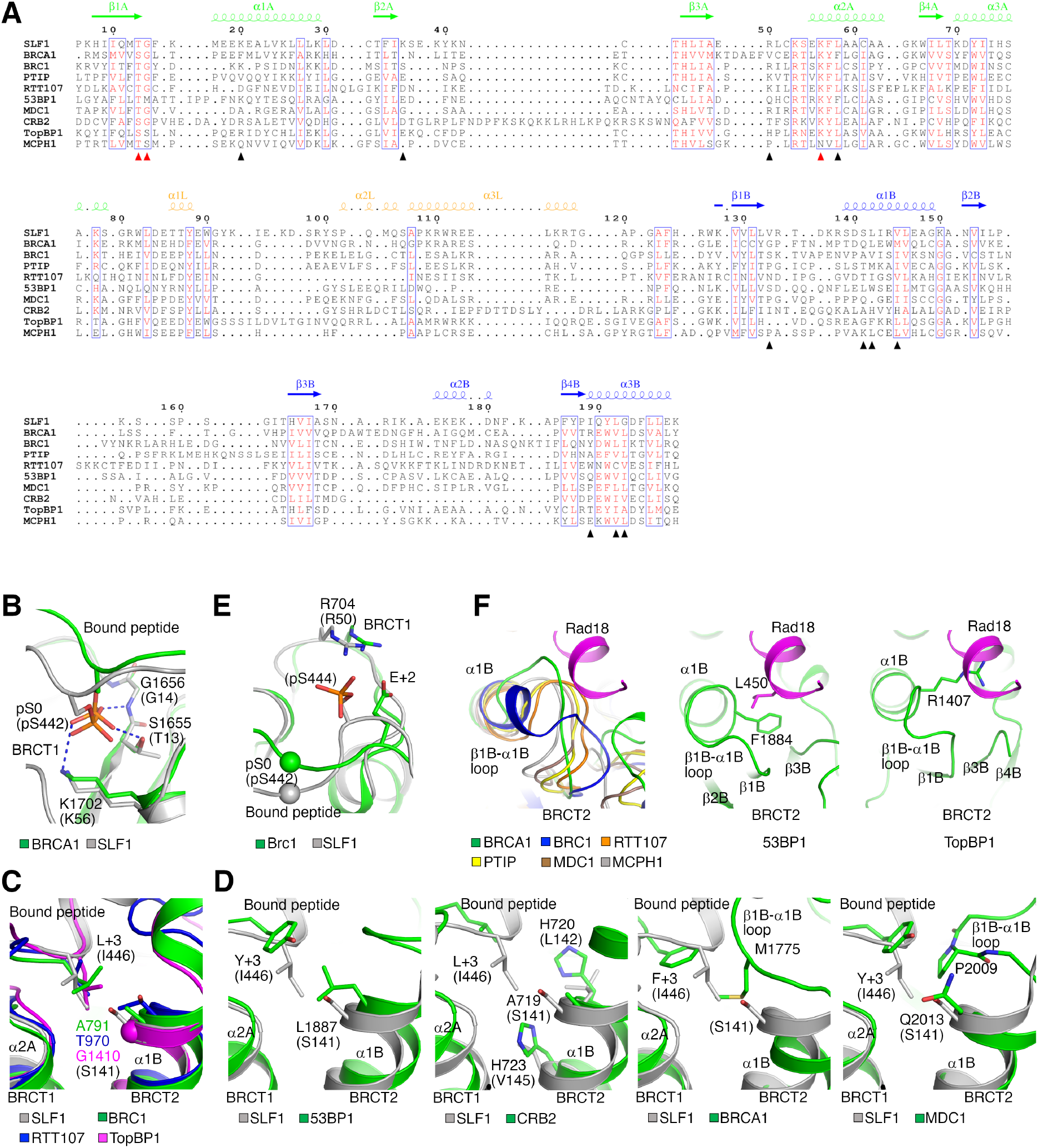
Structural comparison of the SLF1^tBRCT^-Rad18 complex and other tBRCT-ligand complexes. (A) Structure based sequence alignment of SLF1^tBRCT^ and other tBRCTs with similar structures. Residue numbers and secondary structural elements of SLF1 are indicated. The red and black triangles indicate residues interacting with pS442 and additional regions in Rad18, respectively. The alignment is produced by US-align(44). (B) Interactions with pS442 in Rad18 or the anchoring phosphoserine in BRCTide (pS0) bound to the BRCA1 tBRCT (PDB 1T2V). Residue numbers in parentheses are for SLF1 and Rad18. (C) A deep binding pocket for Rad18 Ile446 in SLF1^tBRCT^. Structures of tBRCTs in Brc1 (PDB 3L41), Rtt107 (PDB 3T7K) and TopBP1 (PDB 3AL3) with bound peptides are superimposed for comparison. (D) Shallow binding pockets for the +3 ligand residue in tBRCTs. Structures of tBRCTs in 53BP1 (PDB 5ECG), Crb2 (PDB 2VXC), BRCA1 (PDB 1T2V) and MDC1 (PDB 2AZM) with bound peptides are shown. The structure of the SLF1^tBRCT^-Rad18-2P complex is superimposed for comparison. (E) The interaction between Arg704 in Brc1 and Glu +2 in the bound peptide (PDB 3L41). The structure of the SLF1^tBRCT^-Rad18-2P complex is superimposed for comparison. The spheres indicate pS442 in Rad18 or the anchoring phosphoserine in the Brc1-bound peptide (pS0). (F) Other tBRCTs may not accommodate a Rad18-like helical structure. The β1B-α1B loop and surrounding regions in tBRCTs in BRCA1 (PDB 1T2V), Brc1 (PDB 3L41), Rtt107 (PDB 3T7K), PTIP (PDB 3SQD), MDC1 (PDB 2AZM), MCPH1 (PDB 3SZM), 53BP1 (PDB 5ECG) and TopBP1 (PDB 3AL3) are shown. The Rad18 helix is modeled based on structure alignment of SLF1^tBRCT^ and these tBRCTs. It forms steric clashes with the β1B-α1B loop in these tBRCTs.

The previously characterized tBRCTs contain two common ligand binding pockets. The first located in BRCT 1 is conserved and interacts with the anchoring phosphoserine or phosphothreonine in the ligand peptide, the second is located between the two BRCT domains and recognizes the +3 ligand residue (3 residues C-terminal to the anchoring residue) (16,24-33). Equivalent pockets can be found in SLF1^tBRCT^. The first interacts with pS442 in Rad18 and the interactions are similar to the anchoring residue-mediated interactions in other tBRCT-ligand complexes (Fig. 5B). Residues constituting this binding pocket, Thr13, Gly14 and Lys56, are conserved among tBRCTs (Fig. 5A). The second binding pocket recognizes Ile446 in Rad18. It is relatively deep and structurally similar to the second pocket in Brc1, Rtt107 and TopBP1 (Fig. 5C). Ser141 in SLF1 or the equivalent residues in Brc1, Rtt107 and TopBP1 located at the bottom of the pocket contain small side chains, leading to the formation of a deep pocket. Additional tBRCTs contain a shallow second ligand binding pocket, either due to the large side chains of Leu1887 and H720/H723 in 53BP1 and Crb2, respectively; or the different β 1B-α1B loop conformation in BRCA1, MDC1, PTIP and MCPH1 (Fig. 5D). These observations suggest that tBRCTs utilize several common mechanisms for ligand recognition.

Structural comparison revealed three sets of interactions at the Rad18-SLF1^tBRCT^ interface not previously observed in other tBRCT-ligand complexes. The first involves hydrogen bonds formed with Rad18 residues N-terminal to Ser442 (Fig. 4A). Although several other tBRCT ligand peptides contain multiple residues N-terminal to the anchoring residue (Table S1), they adopt different structures and do not form similar interactions with tBRCTs. The second is the polar interaction between pS444 in Rad18 and Arg50 in SLF1^tBRCT^. Though some other tBRCTs can bind dually phosphorylated ligand peptides(32,34), their interactions with the non-anchoring phosphorylated ligand residue do not resemble the pS444-Arg50 interaction between Rad18 and SLF1. Rather the latter bares partial similarity to the interaction between Arg704 in Brc1 and Glu +2 in the bound H2A tail (Fig. 5E). Equivalent residues in other tBRCT-ligand pairs are located too far apart to form a similar interaction. The third is the hydrophobic interactions mediated by the α-helix in Rad18 and a cleft between α1B, α3B and the β1B-α1B loop in SLF1^tBRCT^. Unlike the Rad18 peptide, all the previously characterized tBRCT-bound ligand peptides adopt extended conformations. In most of the other tBRCTs, the different conformations of the β1B-α1B loop and neighboring regions lead to the formation of a smaller cleft incompatible for α-helix binding (Fig. 5F). In Crb2, the polar side chains of His720 and His723 make the cleft unfavorable for hydrophobic interactions (Fig. 5D). The unique structure of Rad18 presents its Ile446 at the +4 position relative to pS442 for interaction with the second ligand binding pocket in SLF1^tBRCT^. In contrast, the same pocket in other tBRCTs binds the +3 ligand residue. Collectively, these unique interactions at the Rad18-SLF1^tBRCT^ interface contribute to the highly specific Rad18 targeting by SLF1^tBRCT^.

## Discussion

Our structural and biochemical analyses reveal the molecular basis for the Rad18-SLF1^tBRCT^ interaction. We found that Ser442 phosphorylation and an unique α-helix in Rad18 are critical for the Rad18-SLF1^tBRCT^ association, in which Rad18 residues N-terminal to Ser442 and its Ser444 phosphorylation also play important roles. Such quadripartite binding mechanism explains the high affinity and specificity binding between the phosphorylated Rad18 peptide and SLF1^tBRCT^. Structural comparison revealed that while SLF1^tBRCT^ and other tBRCTs utilize two common pockets for ligand recognition, the Rad18-SLF1^tBRCT^ interface also contains additional interactions not found in other tBRCT-ligand complexes. Thus, specific targeting of Rad18 by SLF1^tBRCT^ entails both mechanisms common among tBRCTs and unique mechanisms.

Rad18 residues involved in its recognition by SLF1^tBRCT^ are conserved in higher vertebrates but not in fishes or amphibians (Figs 1A and S1A). In contrast, SLF1 orthologs in most vertebrates contain an N-terminal tBRCT (Fig. S1B). Therefore, the Rad18-SLF1^tBRCT^ interaction probably emerges late in evolution and SLF1^tBRCT^ may possess functions beyond Rad18 targeting. In yeast, the SLF1 homolog Nse5 does not contain the tBRCT domains. However, its partner Nse6 can interact with the tetra-BRCT domain in Rtt107/Brc1, which also contains a tBRCT domain that binds to γH2A at DNA lesions sites. Through interaction with Nse6, Rtt107/Brc1 recruits the SMC5/6 complex to DNA damage sites(17,35-37). Thus, in both yeast and higher eukaryotes, BRCT domains can target the SMC5/6 complex to DNA lesion sites though the exact mechanism varies.

Previous cellular studies indicated that the Rad18 interacts with SLF1 in a phosphorylation dependent manner and the interaction is abolished by the combinatory S442A/S444A substitution in Rad18(5,9). Therefore, the mechanism determined here is likely critical for the interaction between the full length SLF1 and Rad18 *in vivo*. The kinases phosphorylating Ser442 and Ser444 in Rad18 are currently unknown. Future studies to identify these kinases will help to better understand the Rad18-SLF1 signaling cascade. The Rad18 pS444 side chain is solvent exposed in our structure, suggesting that it has the potential for binding additional factors to relay the phosphorylation signal. Future studies are required to test this idea and further define the pathways involving Rad18, SLF1, and the SMC5/6 complex.

## Experimental procedures

### Protein expression and purification

The coding region for SLF1^tBRCT^ (residues 1–199) was optimized for expression in *E. coli*, synthesized (Sangon Biotech) and inserted into the vector pET26b (Novagen). The recombinant protein contains a C-terminal 6x histidine (his-) tag. For protein expression, *E. coli* BL21 Rosetta (DE3) cells transformed with this plasmid were cultured in LB medium supplemented with 34 mg/l kanamycin and 25 mg/l chloramphenicol and induced with 0.25 mM Isopropyl β-D-1-thiogalactopyranoside (Meilunbio) at 16°C for 16 hours. Collected cells were lysed with an AH-2010 homogenizer (ATS Engineering) and SLF1^tBRCT^ was purified by nickel–nitrilotriacetic acid agarose (Smart Life sciences), heparin (Hitrap Heparin HP, GE Healthcare) and gel filtration (Superdex 75 Increase 10/300, GE Healthcare) columns. The protein was concentrated to 10 mg/ml in a buffer containing 20 mM Tris (pH 7.5), 200 mM sodium chloride and 2 mM dithiothreitol (DTT), flash-cooled in liquid nitrogen and stored at -80 °C.

The coding region for the human Rad18 was optimized for expression in *E. coli*, synthesized (Sangon Biotech) and inserted into the vector pET28A (Novagen). Strep-, his- and sumo-tags were introduced to the Rad18 N-terminus by a PCR-based protocol. The recombinant triple tagged Rad18 was expressed in *E. coli* BL21 Rosetta (DE3) cells and purified with streptactin (Smart Life sciences), heparin (Hitrap Heparin HP, GE healthcare) and gel filtration (Superdex 200 10/300, GE healthcare) columns. The protein was concentrated to 10 mg/ml in a buffer containing 20 mM Tris (pH 7.5), 200 mM sodium chloride and 2 mM DTT, flash-cooled in liquid nitrogen and stored at -80 °C.

Amino acid substitutions were introduced with a PCR-based protocol and verified by DNA sequencing. The substituted proteins were expressed and purified following the same protocols for the wild type proteins.

### Crystallization and structure determination

Prior to crystallization experiments, the SLF1^tBRCT^ solution was supplemented with 5x molar excess of Rad18-2P or Rad18-pS442 peptides (Sangon Biotech, Hefei Scierbio-Tech). Crystallization experiments were performed with a sitting-drop setup at 18 °C. The reservoir solution for crystals with the Rad18-2P peptide contains 0.19M magnesium chloride, 0.1M Bis-Tris (pH6.5) and 24% polyethylene glycol (PEG) 3350; for crystals with the Rad18-pS442 peptide contains 0.3M magnesium chloride, 0.1M Bis-Tris (pH6.5) and 24% PEG3350. Prior to diffraction experiments, crystals were equilibrated in the reservoir solution supplemented with 25% (v/v) 2-propanol and flashed cooled in liquid nitrogen. Diffraction data for crystals containing the Rad18-2P peptide was collected at the National Facility for Protein Science beamline BL19U1 at Shanghai Synchrotron Radiation Facility (SSRF) at 0.9785 Å and indexed, integrated and scaled with the HKL3000 package(38). The structure was determined with molecular replacement with PHASER(39), with the structure of the TopBP1 BRCT7/8 repeat (PDB 3AL2) as the search model. Diffraction data for crystals containing the Rad18-pS442 peptide was collected at the SSRF beamline BL02U1 at 0.9792 Å and indexed, integrated and scaled with the XDS package(40). The structure was determined with molecular replacement with PHASER, with the structure of SLF1^tBRCT^ as the search model. Inspection and modification of the structures were carried out with O(41) and COOT(42). The structures were refined with PHENIX(43).

### Isothermal titration calorimetry

ITC experiments were carried out on a MicroCal ITC 200 instrument (Malvern) at 25 °C. Prior to the ITC experiments, SLF1^tBRCT^ was exchanged into a buffer containing 20 mM Tris (pH 7.5) and 200 mM sodium chloride and peptides (Sangon Biotech, Scilight biotechnology, Hefei Scierbio-Tech, Synpeptide) were dissolved in the same buffer. To characterize binding, a solution containing 1.3 mM Rad18 peptide or 6 mM BRCTide was injected into a 300-μl cell that stores 0.1 mM (for binding experiments with Rad18 peptides) or 0.2 mM (for the binding experiment with BRCTide) SLF1^tBRCT^, 2 μl at a time. Data were analyzed with ORIGIN 7.0 (Originlab).

### Pull-down experiments

To characterize Rad18 binding to SLF1^tBRCT^, 50 μg his-strep-sumo-tagged Rad18 was incubated with 30 μl streptactin beads (Smart Life sciences) in a binding buffer containing 20 mM Tris (pH 7.5), 200 mM sodium chloride and 2 mM DTT for 60 minutes at 4 °C. After washing with 800 μl binding buffer three times, the beads were incubated with 50 μg SLF1^tBRCT^ at 4 °C for 60 minutes. The beads were washed again with 800 μl binding buffer three times and the bound proteins were eluted with the binding buffer supplemented with 10 mM D-desthiobiotin (Sigma) and analyzed with SDS PAGE.

### Data availability

Diffraction data and refined structures of SLF1^tBRCT^ in complex with the Rad18-2P or Rad18-pS442 peptides have been deposited into the protein data bank (www.rcsb.org), with accession codes 8IR2 and 8IR4, respectively.

## Acknowledgements

We thank scientists at the National Facility for Protein Science beamline BL19U1 at SSRF and the SSRF beamline BL02U1 for setting up the beamlines and assistance during diffraction data collection, the Large Equipment Sharing Platform at Tianjin Medical University for assistance with ITC experiments.

## Supporting information

This article contains supporting information.

## Conflict of interest

The authors declare no conflict of interest.

## Funding

This work is supported by Natural Science Foundation of China (general grants 32271259 and 32071205 to SX, 32100980 to MS).

## Notes

### Competing Interest Statement

The authors have declared no competing interest.

