## Supplementary material for "Structural insights into Rad18 targeting by the SLF1 BRCT domains": suplemental Figures S1-S5, supplemental Table S1

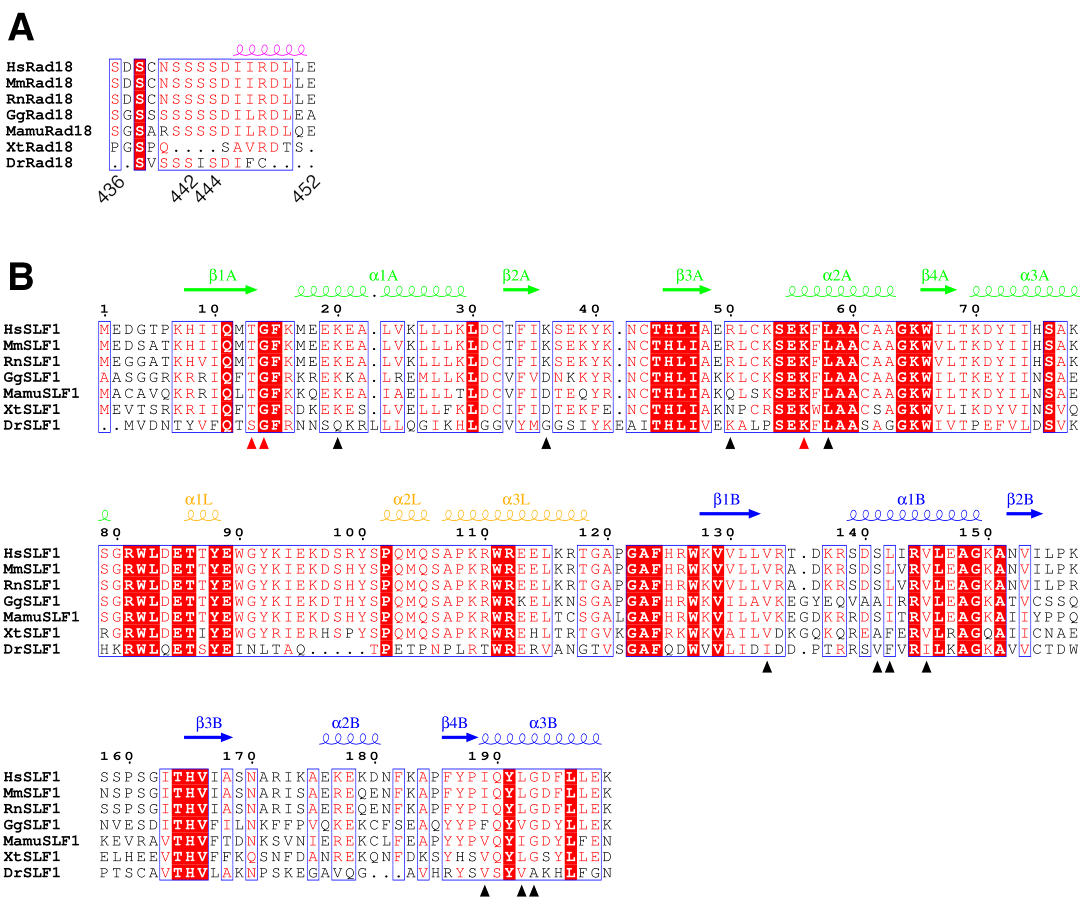

Figure S1 Sequence alignment of Rad18 (A) and SLF1$^{tBRCT}$ (B). Residue numbers and secondary structure elements for the human Rad18 and SLF1$^{tBRCT}$ are shown. The red and black triangles indicate SLF1 residues interacting with pS442 and additional regions in the human Rad18, respectively. Xt, *Xenopus tropicalis* (frog); Dr, *Danio rerio* (zebra fish).

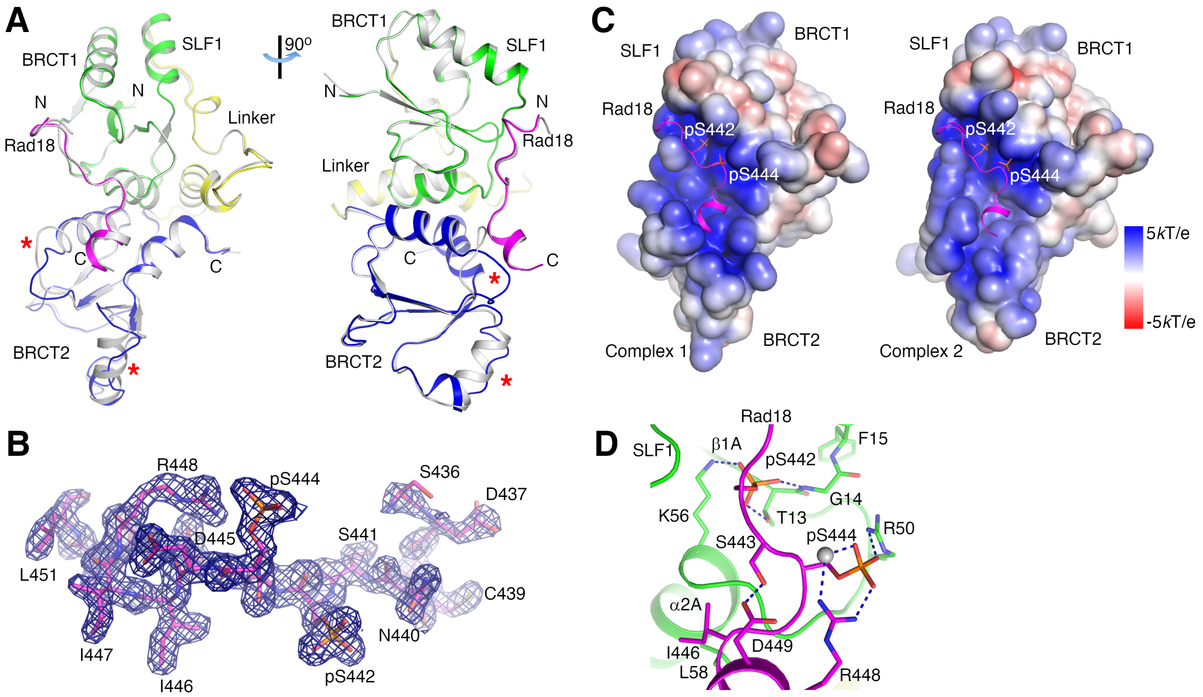


Figure S2 Structure of the SLF1$^{tBRCT}$-Rad18-2P complex. (A) Structure comparison of the two complexes in the asymmetric unit. The second complex is colored in gray. The red stars indicate regions with large structural differences. (B) Electron density map for the second Rad18-2P peptide in the asymmetric unit. The densities are contoured at 1σ. (C) Surface charge distribution of SLF1$^{tBRCT}$. Complexes 1 and 2 in the crystal are shown. Cartoon diagrams of the bound Rad18 peptides are shown for reference. (D) Interactions mediated by pS442, pS444 and surrounding regions in Rad18. Structure of complex 2 in the crystal is presented.

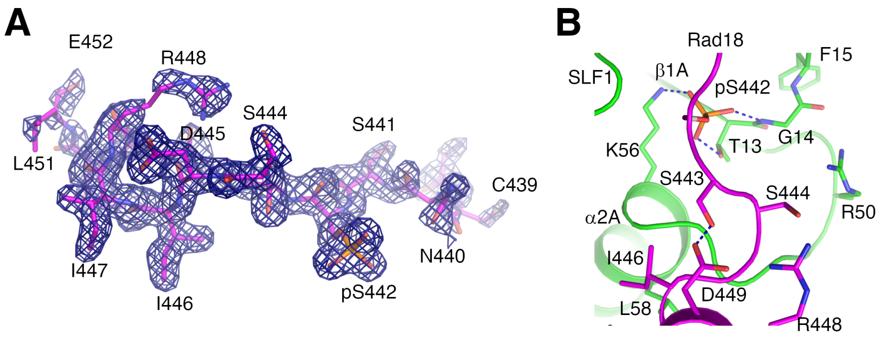


Figure S3 Structure of the SLF1$^{tBRCT}$-Rad18-pS442 complex. (A) Electron densities for the Rad18-pS442 peptide. Electron densities for the Rad18 peptide bound to SLF1$^{tBRCT}$ molecule 2 in the crystal contoured at 1σ are presented. (B) Interactions mediated by pS442 and surrounding regions in the Rad18-pS442 peptide. Structure of complex 2 in the crystal is presented.

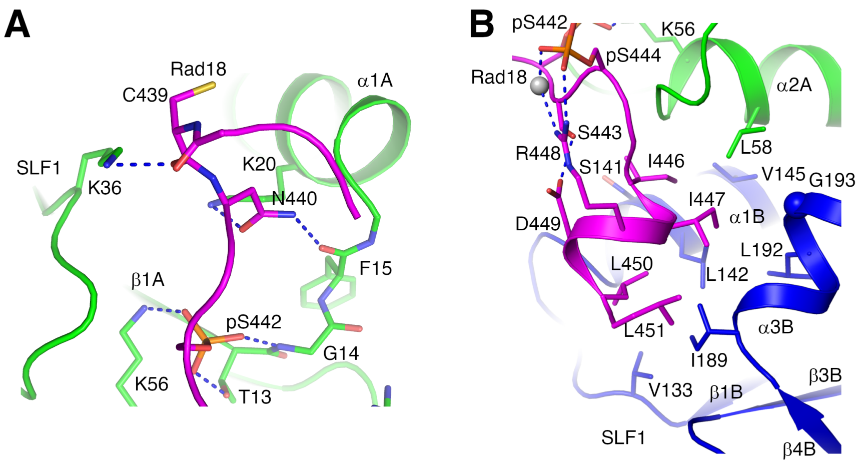


Figure S4 Rad18 regions adjacent to Ser442 and Ser444 contribute to its interaction with $SLF1^{tBRCT}$. Interactions mediated by the N- (A) and C-terminal (B) regions of the Rad18-2P peptide in complex 2 in the crystal are presented.

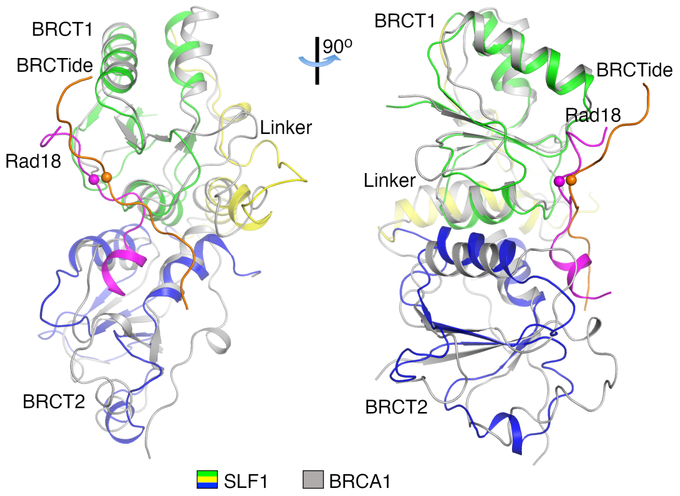


Figure S5 Structural comparison of tBRCTs in SLF1 and BRCA1 (PDB 1T2V, shown in gray). Their structures can be aligned with a RMSD of 1.83 Å for the Cα atoms. The BRCA1-bound BRCTide is shown in brown. The spheres indicate pS442 in Rad18 or the anchoring phosphoserine in BRCTide.

Table S1 Sequences of tBRCT-bound phosphopeptides[1]

| PDB | tBRCT factors | Binding partners | Binding partner peptide sequences[2] |
|---|---|---|---|
| | SLF1 | Rad18 | SDSCNSSSSDIIRDLLE |
| 1T15 | BRCA1 | BACH1 | STSPTFNK |
| 1T29 | BRCA1 | BACH1 | ISRSTSPTFNKQTK |
| 1T2V | BRCA1 | BRCTide | GAAYDISQVFPFAKKK |
| 1Y98 | BRCA1 | CtIP | PTRVSSPVFGAT |
| 3COJ | BRCA1 | Acetyl-CoA Carboxylase 1 | DSPPQSPTFPEAG |
| 3K0H | BRCA1 | Minimal sequence | SPTF-$NH_2$ |
| 3K0K | BRCA1 | Minimal sequence | SPTF |
| 3K15 | BRCA1-D1840T | Minimal sequence | SPTF-$NH_2$ |
| 3K16 | BRCA1-D1840T | Minimal sequence | SPTF |
| 3PXE | BRCA1-E1836K | BACH1 | SRSTSPTFNK |
| 4IFI | BRCA1 | BAAT | RSPVFS |
| 4IGK | BRCA1 | ATRIP | ACSPQFG |
| 4JLU | BRCA1 | Abraxas | GFGEYSRSPTF |
| 4U4A | BRCA1 | Abraxas | GFGEYSRSPTF |
| 4Y18 | BRCA1 | Abraxas | GFGEYSRSPTF |
| 4Y2G | BRCA1 | Abraxas | YSRSPTF |
| 3L41 | BRC1 (BRCT 5/6) | γH2A | KPSQEL |
| 3T7K | Rtt107 (BRCT 5/6) | γH2A | ATKASQEL |
| 3SQD | PTIP (BRCT 5/6) | γH2AX | KKATQASQEY |
| 2VXC | Crb2 | γH2A.1 | SQEL |
| 5ECG | 53BP1 | γH2AX | SQEY |
| 2AZM | MDC1 | γH2AX | KKATQASQEY |
| 3K05 | MDC1-T2067D | Minimal sequence | SQEY-$NH_2$ |
| 3SHV | MCPH1 | γH2AX | KKATQASQEY |
| 3SZM | MCPH1 | γH2AX | KKATQASQEY |
| 3U3Z | MCPH1 | γH2AX | SQEY |
| 3T1N | MCPH1 | Cdc27 | SDEF |
| 3AL3 | TopBP1 (BRCT 7/8) | BACH1 | SIYFTPELYD |

[1]Sequences of peptides bound to tBRCTs structurally homologous to SLF1$^{tBRCT}$ are listed.
[2]Phosphorylated residues are colored in red, disordered residues in the structure are colored in gray. The anchoring phosphoserine or phosphothreonine are aligned.
